# Adaptive spatio-temporal phase retrieval for ultra-short pulse wavefront shaping

**DOI:** 10.64898/2026.02.04.703770

**Authors:** Oz Shaul, Tali Ilovitsh

## Abstract

Beam shaping of ultra-short pulses is essential for medical ultrasound, where single-cycle excitations are required to achieve high axial resolution and improve frame rate. Conventional methods, such as the Gerchberg–Saxton (GS) algorithm or more recent deep learning approaches, are generally effective for continuous-wave excitation but degrade significantly under single-cycle conditions. In diagnostic imaging, high frame rate is critical for applications demanding rapid scanning. In this context, multi-line transmission (MLT) leverages beam shaping to synthesize multiple simultaneous foci, thereby increasing frame rate. In parallel, structured illumination methods for super-resolution and acoustical holography likewise depend on actively shaping single-cycle pulses to produce controlled patterns, highlighting the need for precise short-pulse beam shaping. To address this challenge, we introduce the spatio-temporal adaptive reconstruction (STAR) algorithm, which performs active beam shaping directly in the time domain by integrating the generalized angular spectrum method (GASM) into an iterative optimization scheme. STAR enforces constraints on both the transducer and focal planes, enabling accurate control of single-cycle excitations. Simulations showed that STAR consistently outperformed GS for multi-focus patterns. For example, in a four-foci configuration, STAR achieved a correlation of 0.80 compared to 0.64 for GS, with significantly improved uniformity across focal peaks. Resolution analysis demonstrated that STAR reduced the minimum distinguishable foci spacing from 1.09 mm with GS to 0.87 mm. Experimental hydrophone measurements confirmed these improvements. Across multi-foci patterns, STAR produced more distinct and balanced foci compared to those observed with GS. These results demonstrate that STAR provides robust and efficient active beam shaping of single-cycle pulses, maintaining accuracy across different depths and frequencies for diagnostic applications.

## I INTRODUCTION

Wavefront shaping is a fundamental concept in various wave-based technologies, including optics, acoustics, microwaves, and biomedical imaging. It enables precise control over the spatial and temporal properties of waves, allowing tailored energy distribution for specific applications. Applications such as high-resolution imaging [1], optical trapping [2],[3], focused energy delivery [4] and acoustic holography [5] rely on the ability to manipulate wavefronts to achieve desired field distributions. Various methods have been developed to accomplish this, ranging from iterative phase-retrieval algorithms to optimization-based techniques and, more recently, deep learning-driven approaches.

One of the most established classes of wavefront shaping techniques is iterative phase-retrieval methods, where an initial wavefront is iteratively refined to match a target intensity distribution. The Gerchberg-Saxton (GS) algorithm [6], one of the most popular methods in this category, alternates between the transducer plane and the focal plane, enforcing constraints in each domain to converge toward the desired beam profile. Variants of the GS algorithm introduce refinements to enhance accuracy, convergence speed and robustness [7],[8],[9]. Another widely used category is optimization-based wavefront shaping, where numerical techniques, such as gradient descent [10], genetic algorithms [11] and automatic differentiation [12] are employed to iteratively refine the input wavefront. These methods offer higher accuracy but are often computationally expensive, limiting their applicability in real-time settings. More recently, deep learning-based approaches have been explored for wavefront shaping. These include convolutional neural networks (CNNs) [13],[14],[15], autoencoder frameworks [16],[17], and generative adversarial networks (GANs) [18], which enable direct mapping between input and desired wavefronts. These methods have demonstrated improved performance in complex scenarios but often require large training datasets and lack interpretability. Among these approaches, the GS algorithm remains widely used due to its simplicity, rapid convergence and flexibility in optimizing different parameters such as frequency and depth. However, while effective for continuous-wave (CW) signals, the GS algorithm, and similar methods, struggle with ultra-short pulses, particularly single or two-cycle pulses. This limitation stems from the reliance on wave propagation models which assume steady-state or CW insonation, such as the angular spectrum method (ASM).

To overcome these limitations, we propose the spatio-temporal adaptive reconstruction (STAR) algorithm that enables direct time-domain optimization for precise wavefront shaping of ultra-short pulses. Ultra-short pulses are critical across multiple fields, as they enable improved resolution, and reduced interference outside the region of interest. In radar and wireless communications, ultra-short pulses provide better resolution, precise distance measurement, improved target recognition, and enhanced resistance to noise and interference [19]. In optical microscopy and super-resolution imaging, techniques such as two-photon fluorescence microscopy [20] and stimulated emission depletion (STED) [21] microscopy rely on ultra-short laser pulses to achieve deeper penetration depths and sub-diffraction-limited resolution. In medical ultrasound imaging, ultra-short pulses play a crucial role in achieving high axial resolution, enabling the distinction of closely spaced anatomical structures [22].

High frame rate is particularly demanded for broad range of imaging applications such as echocardiography and blood flow imaging, where capturing rapid physiological dynamics is essential. One investigated strategy to achieve high frame rate imaging is multi-line transmission (MLT), in which multiple focused beams are transmitted simultaneously. This approach allows significant gains in frame rate without strongly compromising spatial resolution or signal-to-noise ratio [23],[24],[25],[26]. However, MLT performance critically depends on the ability to shape beams accurately under single-cycle pulse excitation. Additional applications include structured illumination super resolution methods [1],[27], where there is a need to generate multiple simultaneous foci with high precision.

To address the need for wavefront shaping with ultra-short pulses, a deep convolutional residual network-based approach was developed [28]. This study demonstrated superior performance compared to the GS algorithm in terms of accuracy and computational efficiency. However, because the training dataset spans limited frequency and depth ranges, generalization is limited, and parameter changes require retraining. This study aims to develop a wavefront shaping algorithm that retains the adaptability of the GS algorithm while enabling robust performance for ultra-short pulses, including single-cycle excitations and other transient signal types.

A key technique for modeling wave propagation in ultrasound and other wave-based applications is the ASM [29], which models wavefields by decomposing them into plane waves with distinct propagation directions and amplitudes. Initially designed for CW signals, ASM has been instrumental in ultrasound applications due to its computational efficiency and accuracy, making it a core component of the GS algorithm as well. However, the conventional ASM struggles with short pulses due to its inherent reliance on steady-state, harmonic solutions [30]. The broadband, transient nature of short pulses necessitates a generalized approach to adequately capture their propagation dynamics. To address this limitation, the ASM has been extended to accommodate pulsed ultrasound signals, leveraging a time-domain framework to account for the transient nature of these signals [31]. This generalized method, generalized angular spectrum method (GASM), provides a robust foundation for modeling the propagation of short pulses, enabling accurate simulation of acoustic fields for advanced imaging applications.

Several approaches have been proposed for reconstructing nonstationary acoustic fields directly in the time domain [32],[33],[34],[35]. These methods enable reconstruction of transmitted waveforms without assuming continuous-wave excitation. However, they remain passive reconstruction techniques: they do not impose physical constraints on the transmitted signals or the desired pattern and therefore cannot optimize the waveforms toward forming a prescribed target field. Moreover, while the above studies have been demonstrated for sound fields, they have not addressed higher-frequency ultrasound.

Here, we propose the STAR algorithm which provides a framework for transmission-side, time-domain active beam shaping of ultra-short pulses. By integrating the GASM into the GS framework, the STAR algorithm overcomes the constraints of phase-domain optimization, allowing for precise wavefront shaping of ultra-short pulses.

## II Methods

### A Angular Spectrum Method

Mathematically, the ASM can be expressed as follows. Let the initial pressure field at the transducer surface be given by *p*(*x,y,z*_0_). The angular spectrum of this field is defined as the two-dimensional Fourier transform:

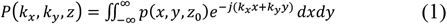

where *k*_*x*_ and *k*_*y*_are the spatial frequency components in the *x* and *y* directions, respectively, and *z* is the position of the given plane in the z direction.

The pressure field at a plane in depth of *z*_1_ can then be calculated as:

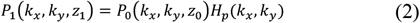

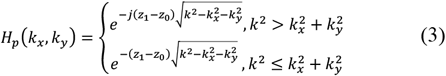

where *p*_0_ and *p*_1_ are the angular spectrums in depths of *z*_0_ and *z*_1_ respectively, and *k* is the wavenumber, with *f* being the frequency and *c* the speed of sound in the medium.

### B Angular Spectrum Method for Pulsed Fields

This method can be generalized via GASM to calculate the pressure field for multi-frequency signal, as follows:

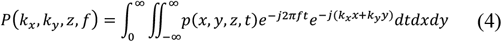

where now *p*(*x, y, z, t*) is now dependent on time and *p*(*k*_*x*_, *k*_*y*_, *z, f*)is the angular spectrum which is dependent also on the temporal frequency, *f*, as the 3D fourier transform was calculated (over x,y and t).

The relation between the angular spectrums in parallel planes (in depths of *z*_0_ and *z*_1_) is:

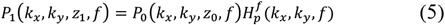

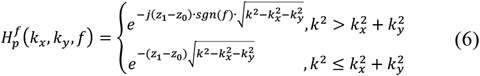

And finally the pressure field in depth of *z*_1_is given by:

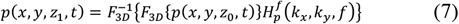

This generalized formulation allows for the accurate modeling of the propagation of short, pulsed signals in general and specifically in ultrasound imaging, as it takes into account the temporal evolution of the pressure field, in addition to the spatial distribution.

### C. Simulations

Both methods were implemented in MATLAB and demonstrated by simulating signal propagation from a P6-3 transducer (Philips, Bothell, WA, USA), that was used in the experiments. The P6-3 transducer features 128 elements, a center frequency of 4.46 MHz, and a pitch of 0.22 mm.

Validation of both methods was performed through comparison with the k-Wave Simulation Tool, a robust numerical framework for modeling acoustic wave propagation. Unlike the previously discussed analytical methods, k-Wave utilizes a computational, grid-based approach to simulate the propagation of acoustic waves.

The ASM was validated against k-Wave simulations implemented in MATLAB by transmitting long pulses exceeding 10 cycles (more than 2 µs) in duration. The GASM, on the other hand, was validated using 1-cycle pulses (∼0.22 µs) for a direct comparison with the proposed approach.

The toneBurst function from the k-Wave toolbox was used to generate ultrasound signals for each transducer element during implementation of both the GASM and the STAR algorithm presented subsequently.

### D Gerchberg-Saxton Algorithm

The GS algorithm can be summarized as follows (Alg. 1.): First, the phase distribution across the transducer elements is initialized to be uniform. Next, the initial pressure field is propagated to the focal plane using the ASM, with the target depth set to 40 mm. At the focal plane, the desired amplitude constraint is applied while preserving the phase information. Subsequently, inverse propagation is calculated to obtain the updated pressure field in the transducer plane. An amplitude constraint is then applied in the transducer plane by setting the amplitude to 1 for x values within the transducer’s region and 0 for those outside it. These steps are repeated iteratively until the algorithm converges to the desired beam profile.

After the final iteration of the GS algorithm, the resulting phase distribution across the transducer elements is unwrapped. The motivation for this unwrapping step is to remove any discontinuities in the phase and better exploit the time-domain properties of the signal. The unwrapping process involves adding or subtracting multiples of 2π to the phase values, ensuring a smooth, continuous phase function.

Then, the resulting apodization-phase distribution across the transducer elements can be used to drive the ultrasound system and generate the desired acoustic beam shape.

### E. STAR Algorithm for Pulsed Signal

The proposed STAR algorithm beam shapes the acoustic field by iteratively propagating the spatiotemporal signal waveform between the transducer plane and the focal plane using the GASM propagation function, while enforcing constraints on amplitudes, delays, and pulse duration in both spatial and temporal domains. The algorithm utilizes 1-cycle excitations for each transducer element. The STAR algorithm can be summarized as follows (steps are numbered in Alg. 2):

#### Algorithm 1

Gerchberg-Saxton Algorithm

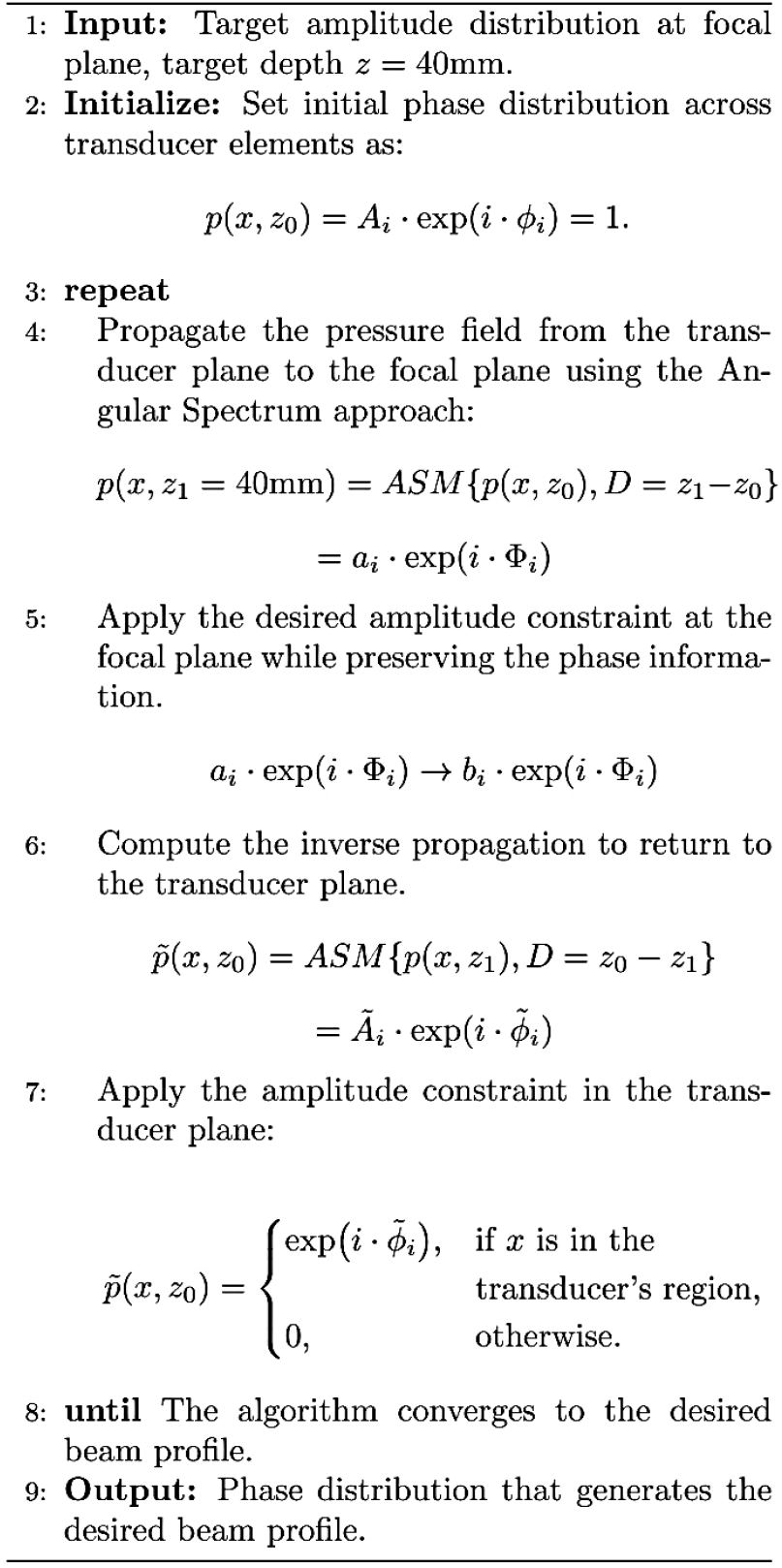

Signals for each transducer element are initialized as single-cycle pulses with zero delay and uniform apodization (step 2). The initial pressure field is then propagated to the focal plane at the target depth of 40 mm using the GASM (step 4). Amplitude constraints are applied in the focal plane to match the desired beam pattern, while the original signal shape and temporal delays are preserved (step 5). The updated pressure field is obtained by performing inverse propagation back to the transducer plane (step 6), where the maximum peak pressure and its temporal location for each transducer element are determined (step 7). Signal duration constraints are applied at the transducer plane, ensuring that each element’s signal remains a single-cycle pulse. The temporal peak locations identified earlier are used as reference delays, and unity apodization is implemented within the transducer region to maximize transmitted energy, while the amplitude is set to zero outside this region (step 8). These steps are iteratively repeated until convergence to the desired beam pattern is achieved. The output pattern is obtained by taking the maximum peak value of the real part of each signal in the focal plane after step 4 of the algorithm. This provides the amplitude distribution across the spatial domain at the target depth.

#### Algorithm 2

STAR Algorithm

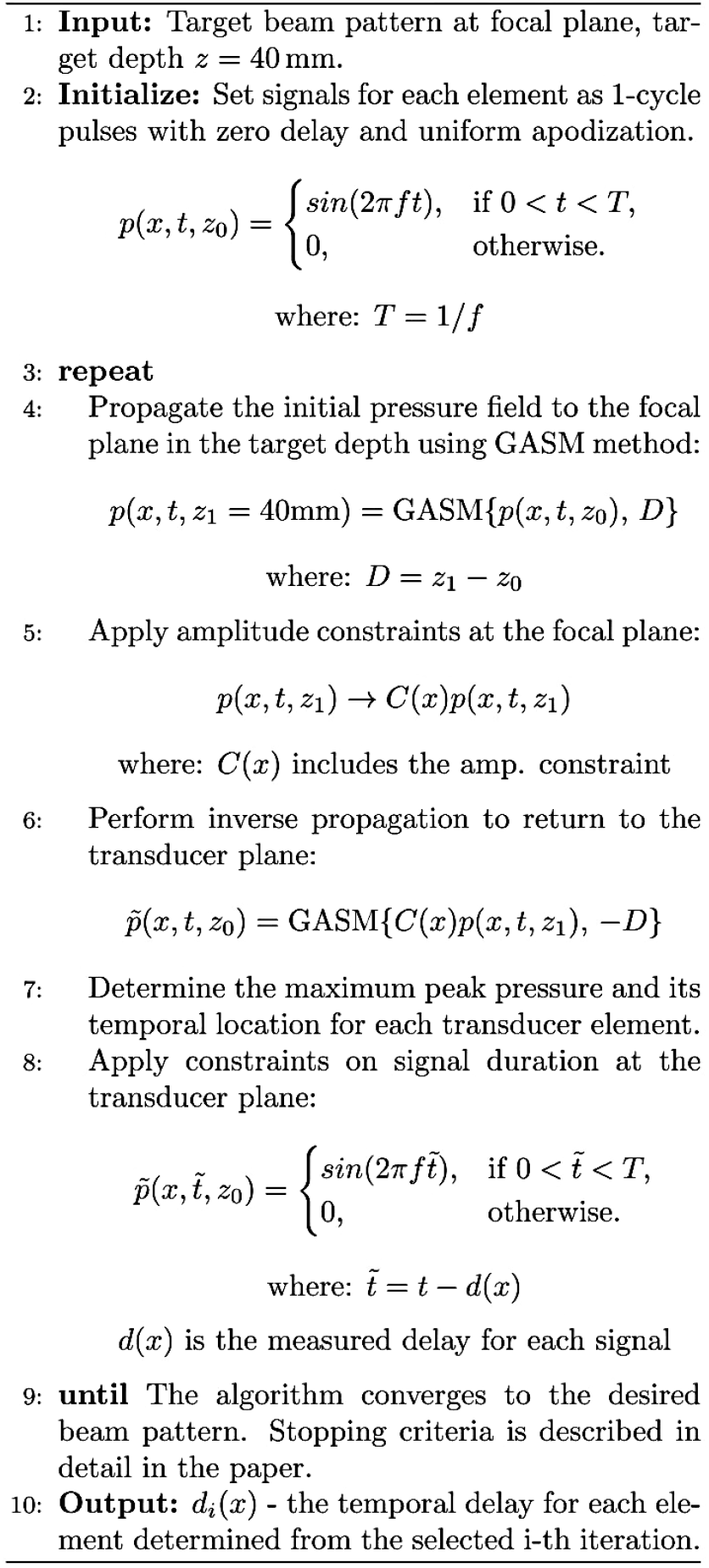

A schematic illustration of the algorithm is provided in Fig 1. Two key parameters are used to measure the success in designing the desired pattern: 1. Normalized cross-correlation between the desired pattern and the output pattern: This value should be maximized, as we want the output pattern to match the desired pattern as closely as possible. However, the maximum achievable value is 1, as the patterns cannot be identical. 2. Peaks Uniformity Coefficient: This metric quantifies the uniformity of the focal heights across the focal plane. It is calculated as the ratio of the standard deviation to the mean of the focal heights. A lower value indicates more similar peak amplitudes, which is desirable for applications like enhancing super resolution. By optimizing these two parameters through the iterative STAR algorithm, the goal is to generate an output pattern that closely matches the desired target while maintaining uniformity in the focal plane.

**Fig. 1.**
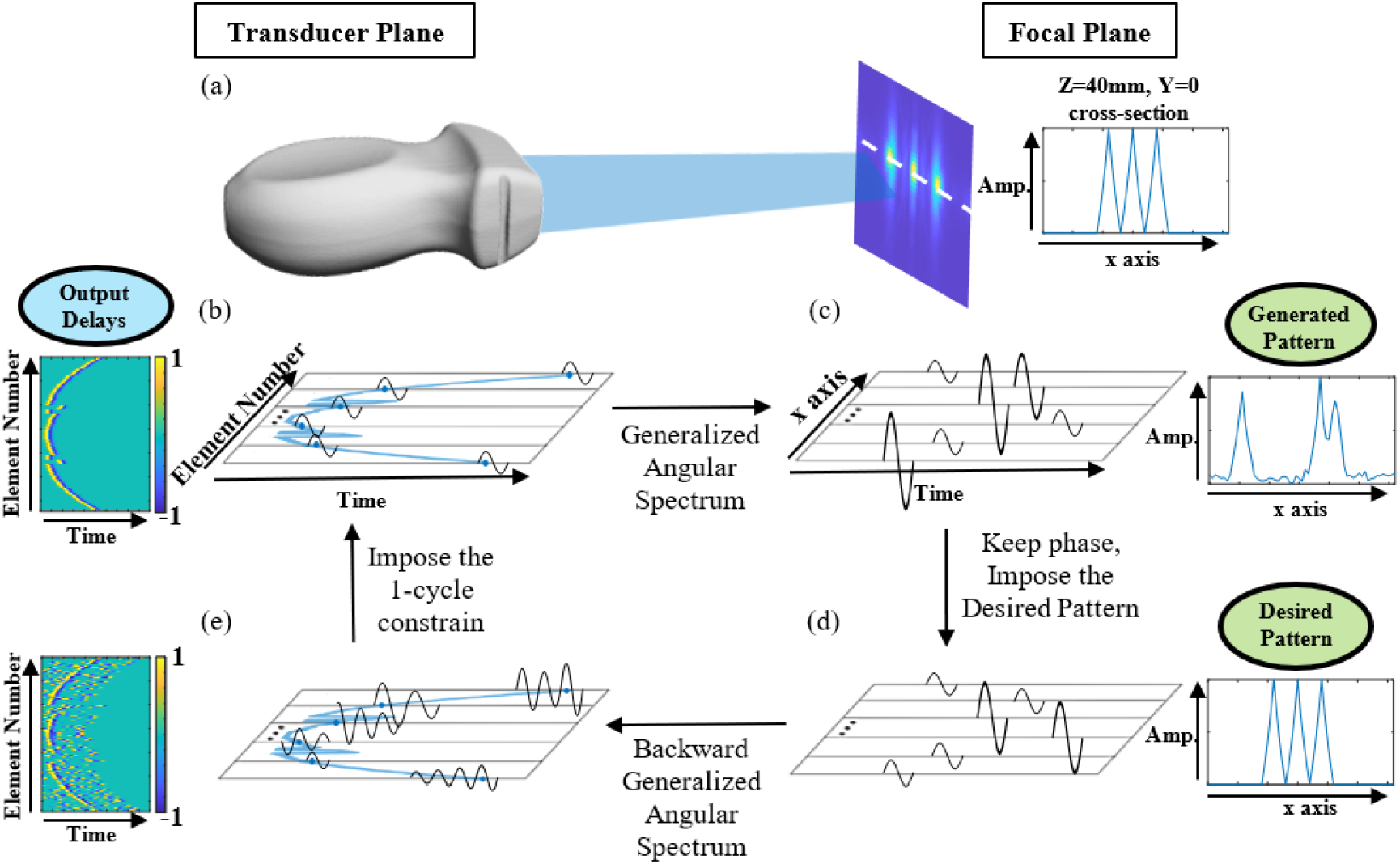
Schematic illustration of the STAR algorithm. (a) A transducer emitting the acoustic field which propagates to the focal plane. At the focal plane, the generated acoustic pattern at the target depth is displayed along with its cross-section, providing a visual reference for the expected algorithm output. (b) Initial single-cycle signals, identical for all elements in the first iteration, are transmitted and propagated forward using the GASM. (c) The propagated signals at the focal plane exhibit varying amplitudes and delays, from which the output pattern is extracted by identifying the maximum peak pressure at each element. (d) The application of the desired pattern constraint, where delays remain unchanged while amplitudes are normalized according to the target pattern. Because the desired pattern is primarily concentrated along the center of the x-axis, elements in the middle exhibit higher amplitudes than those at the edges. (e) The signals are backward-propagated to the transducer plane, where they are no longer single-cycle. Next to this, the signals are represented as a colormap, providing a visualization of their structure. Finally (back in (b)), the transducer plane constraint is enforced, thus signals are truncated to single-cycle duration with uniform amplitude, and the delays are derived from the maximum peak locations, forming the final output of the algorithm.

The algorithm’s output is the temporal delays for each transducer element, determined from the selected iteration. The stopping criteria considers two conditions. First, the Peaks uniformity coefficient must be less than a defined threshold. Second, the normalized cross-correlation should not vary by more than 10% across the last 10 consecutive iterations. If these conditions are not satisfied, the algorithm terminates after 100 iterations and selects the output corresponding to the iteration with the minimum Peaks uniformity coefficient.

Unlike phase-based methods, this approach directly computes the temporal delays, eliminating the need for phase unwrapping. The resulting delay distribution across the transducer elements can then be implemented to shape the desired acoustic beam in the ultrasound system.

### F. Experimental Setup

To validate the proposed algorithm, experimental measurements were performed using a Verasonics ultrasound system (Vantage 256, Verasonics, Kirkland, WA, USA) equipped with a P6-3 phased array transducer. The transducer was programmed to transmit signals from each individual element, with delays calculated using both the STAR and GS algorithms. The transducer emitted signals into a degassed water tank, and the resulting acoustic pressure field was measured using a needle hydrophone (NH0200, Precision Acoustics, Dorchester, U.K.) with an active aperture of 0.2 mm. The hydrophone was mounted on a motorized xyz stage that was controlled by a motion controller (ESP300, Newport, USA). The pressure signals detected by the hydrophone were sampled using a digital oscilloscope (MDO3024, Tektronix, Beaverton, OR, USA) and transferred to a computer for post-processing. The post-processing included generating acoustic pressure maps for the analyzed region. The acoustic field patterns generated by the STAR algorithm were compared to those obtained from simulations as well as to patterns produced using the GS algorithm. For a fair comparison, the transmitted signals for both algorithms were set to a duration of one cycle, consistent with the parameters used in the STAR algorithm. Peak Negative Pressure (PNP) was determined from the measured voltage signal. The PNP at each focus reached up to 0.54 MPa, corresponding to a maximum mechanical index (MI) of 0.25. Notably, as the number of foci increased, the MI decreased accordingly.

## III Results

### A. Validation of GASM and the STAR Algorithm

To validate the quality of the propagation achieved using the GASM for pulses, a comparative analysis was conducted with the propagation performed using the k-Wave toolbox. The comparison was carried out across various patterns and at multiple depths. The transmitted delays for each pattern were computed using either the STAR or the GS algorithms. These algorithms were used to generate patterns with both well-separated foci and less-distinguishable foci to assess performance under different conditions. As a representative example, a 3-foci pattern was evaluated. The delays were computed, and the propagation was performed using both the GASM and the k-Wave toolbox to generate the pattern at a depth of 40 mm. The analysis was conducted over depths ranging from 25 mm to 55 mm relative to the transducer position. Three distinct foci, approximately 6 mm apart, were produced by both approaches and were symmetrically distributed about the center of the x-axis (Fig. 2(a) and Fig. 2(b)). To quantify the differences between the two methods, the mean relative error was calculated at various depths by comparing cross-sections of the propagation. At 40 mm depth, as shown in Fig. 2(d), the cross-sections nearly overlap, with a mean relative error of 1.49%. Across all investigated depths for this pattern, an average relative error of approximately 2% was observed. Fig. 2(c) provides a visual representation of these differences between the XZ-planes generated by GASM and k-Wave. A total of 100 distinct patterns across 300 depth levels (n = 30,000) were analyzed, resulting in a mean relative error of 4 ± 1.5%. After implementing the propagation methods and the GS algorithm, its output was validated. The simplest pattern to test was a single focus at a specific depth, as it allows for direct comparison with an analytical geometrical solution for the delays, given by the negative of the distance of each element divided by the speed of sound. The wavefronts obtained using the algorithm with an input pattern of one focus at the center of the lateral axis, at a depth of 40 mm and a frequency of 4.46 MHz, are shown in Supplementary Fig. 1(a). The output delays match those derived from the analytical geometrical solution, as confirmed by the XZ plane computed through propagation, which accurately reproduces the desired pattern (Supplementary Fig. 1(d)) and its cross-section (Supplementary Fig. 1(g)). As expected, applying these delays to a single-cycle excitation also generates the desired pattern, as shown in the second column of Supplementary Fig. 1. Subsequently, the STAR algorithm was tested using the same pattern and produced identical results, as shown in the right column of the figure. This confirms that the STAR algorithm converges to both the GS algorithm and the analytical solution for the simple yet fundamental pattern of a single focus at the target depth.

**Fig. 2.**
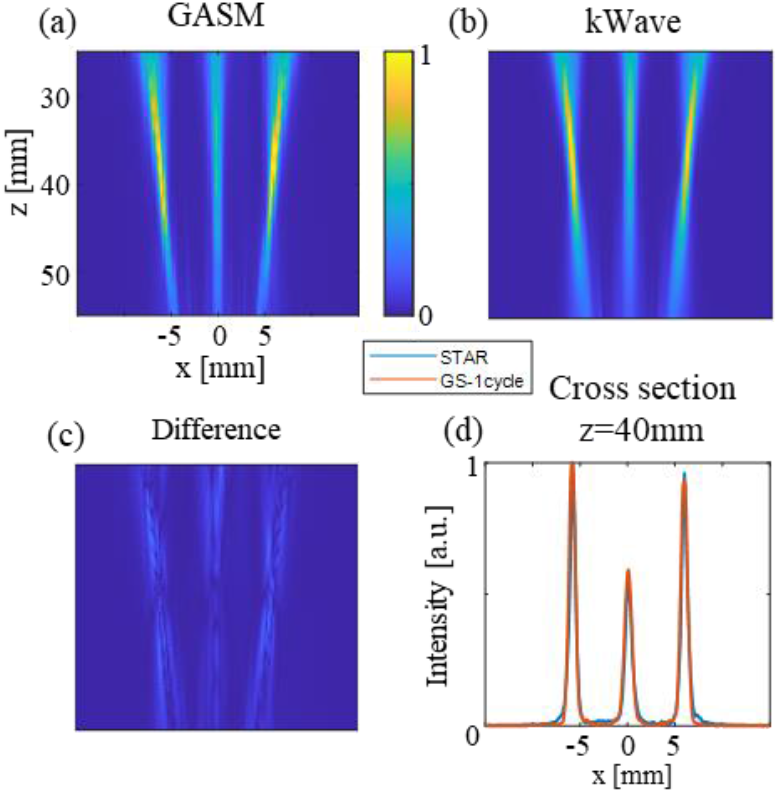
Comparison of pulse propagation using GASM and the k-Wave toolbox for a 3-foci pattern at a depth of 40 mm. The foci are spaced approximately 6 mm apart and symmetrically distributed along the x-axis. (a) Normalized intensity distribution in the XZ-plane obtained using GASM, where X represents the lateral direction and Z the axial direction. (b) Corresponding XZ-plane distribution using the k-Wave toolbox. (c) Visual representation of differences between the XZ-plane results from GASM and k-Wave. (d) Cross-section comparison at 40 mm depth.

### B. Comparison of the GS and STAR Algorithms for a Simple Pattern

After validating the propagation methods, the implementation and convergence of the GS algorithm to the geometrical solution, and the convergence of the STAR algorithm to the GS algorithm in the simple case of a single focus, an example is presented comparing the output of the STAR algorithm to the conventional GS algorithm (Fig. 3). In this example, both algorithms were tasked with generating a 3-foci pattern at a depth of 40 mm, with a spacing of 19 pitches (∼4.15 mm) between each pair of foci, centered around the x-axis. To examine the differences between the methods and the impact of signal duration, signal propagation was computed using the GS output delays for both continuous-wave (CW) and single-cycle excitation, as well as using the STAR output delays for single-cycle excitation. The transmitted signals generated from the phases output by the GS algorithm are illustrated by the signal amplitudes over time for each element (Fig. 3(a)). In this case, with CW excitation, the signals are periodic and consist of multiple cycles. The corresponding acoustic field propagation for this signal is shown with the target depth marked by a white line (Fig. 3(d)). The algorithm achieves accurate results in terms of the location and amplitude of the desired foci, producing three distinct foci precisely at the target depth. However, when using a single-cycle excitation (Fig. 3(b)), the propagated acoustic field (Fig. 3(e)) exhibits less well-defined foci. While three foci are present at the desired locations, two undesired sidelobes appear between them. In contrast, the transmitted signals and propagated acoustic field based on the STAR algorithm delays are shown in Fig. 3(c) and 3(f), respectively. Both Fig. 3(e) and 3(f) are normalized to the same scale to enable a fair comparison, as the emitted energy is identical in both cases due to the single-cycle excitation.

**Fig. 3.**
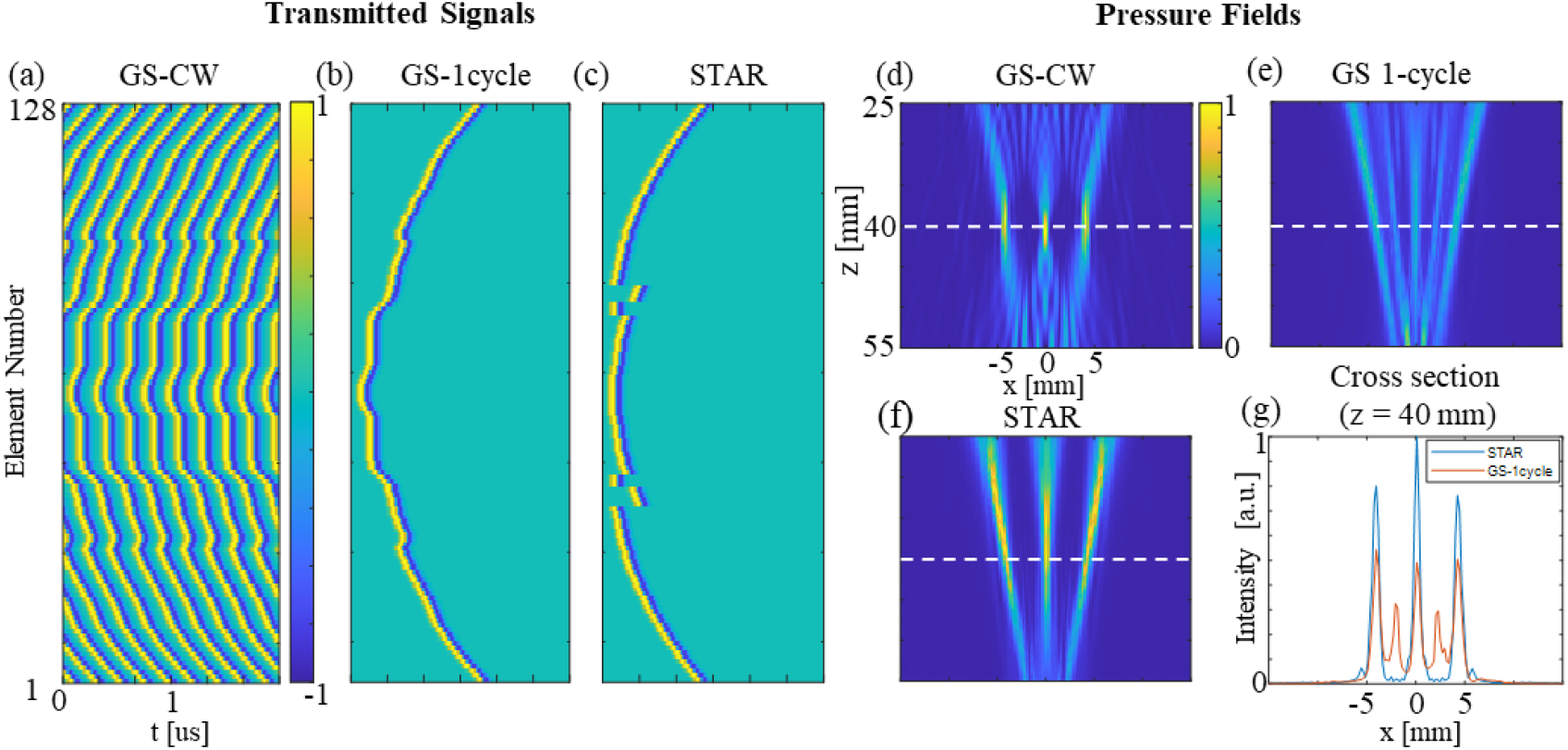
Comparison of the GS and STAR algorithms for generating a 3-foci pattern at 40 mm depth. (a) Transmitted signals for GS output delays with continuous-wave (CW) excitation, with color indicating signal amplitude. (b) Transmitted signals for GS output delays with single-cycle excitation. (c) Transmitted signals for STAR output delays with single-cycle excitation. (d) Propagated acoustic field for GS with CW excitation, with white lines marking the target depth. (e) Propagated acoustic field for GS with single-cycle excitation. (f) Propagated acoustic field for STAR with single-cycle excitation. (g) Cross-section comparisons at 40 mm.

The propagated field from the STAR output was similar to the desired pattern, with three clearly distinguished foci located precisely at the target depth and with heights closely matching the desired values. The cross-sections at the target depth highlight the superior energy concentration at 40 mm achieved with the STAR algorithm compared to the GS algorithm (Fig. 3(g)). The STAR output more closely resembles the desired pattern, effectively demonstrating its advantages in single-cycle excitation scenarios.

### C. Investigation of Key Parameters

While implementing the STAR algorithm, the evolution of key parameters over multiple iterations was investigated. Two primary metrics were analyzed to track their changes across iterations in different examples. An illustrative example is provided using a desired pattern of three foci (Fig. 4). The normalized cross-correlation between the desired and output patterns was examined as a function of the number of iterations (Fig. 4(b)). The GS algorithm converges within a small number of iterations. This behavior is evident, along with a decrease in correlation when switching from CW to single-cycle excitation. However, the STAR algorithm consistently achieves higher correlation values for the same task starting from the second iteration (Fig. 4(b)). Across all cases, the correlation stabilizes after the initial iterations and does not vary significantly thereafter. In contrast, the Peaks Uniformity Coefficient exhibits substantial variability throughout the iterations. A higher number of iterations does not necessarily lead to better uniformity, even if the correlation improves (Fig. 4(e)). This trend is further demonstrated by comparing acoustic fields computed at two different iterations of the STAR algorithm (Fig. 4(a) vs. Fig. 4(c)). By comparing the cross-sections at the target depth, it becomes apparent that the STAR output after 5 iterations better matches the desired pattern in terms of uniform peak heights, despite the correlation being slightly higher after 20 iterations (Fig. 4(d) and 4(f)). This behavior was consistently observed across different patterns, leading to the incorporation of the Peaks Uniformity Coefficient as an additional factor in the stopping criteria and in selecting the specific iteration for output generation.

**Fig. 4.**
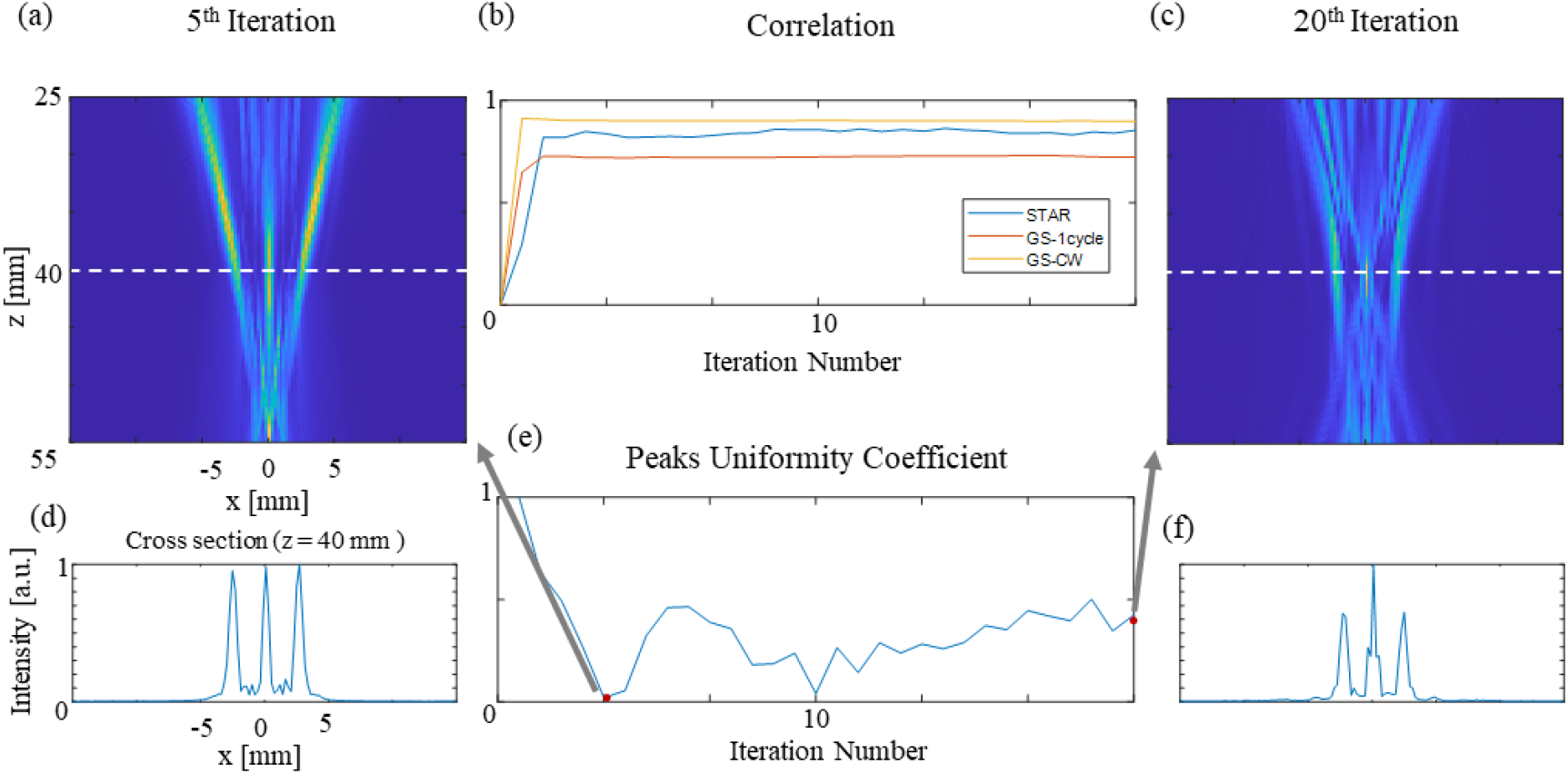
Evolution of key parameters during STAR algorithm iterations for a 3-foci pattern. (a) Output acoustic field after 5 iterations. (b) Normalized cross-correlation between the desired and output patterns as a function of iteration number: GS with CW excitation (yellow), GS with single-cycle excitation (orange), and STAR with single-cycle excitation (blue). (c) Output acoustic field after 20 iterations. (d) Cross-section at 40 mm depth for the STAR output after 5 iterations. (e) Peaks Uniformity Coefficient over iterations. (f) Cross-section at 40 mm depth for the STAR output after 20 iterations.

### D. Performance Comparison of GS and STAR Across Patterns, Depths and Frequencies

The STAR algorithm was tested across various patterns, including configurations with different numbers of foci, varying spacing between adjacent foci, and symmetric or asymmetric arrangements around the lateral axis. The main findings are illustrated through three representative examples (Fig. 5). Each row in Fig. 5 corresponds to a different desired pattern, while each column represents a specific algorithm or excitation type. The first pattern involved two foci spaced approximately 1.5 mm apart, with an offset of about 6 mm from the center of the lateral axis. Both GS and STAR algorithms successfully generated this pattern, concentrating similar amounts of energy at the target depth (Fig. 5(b), 5(c)).

**Fig. 5.**
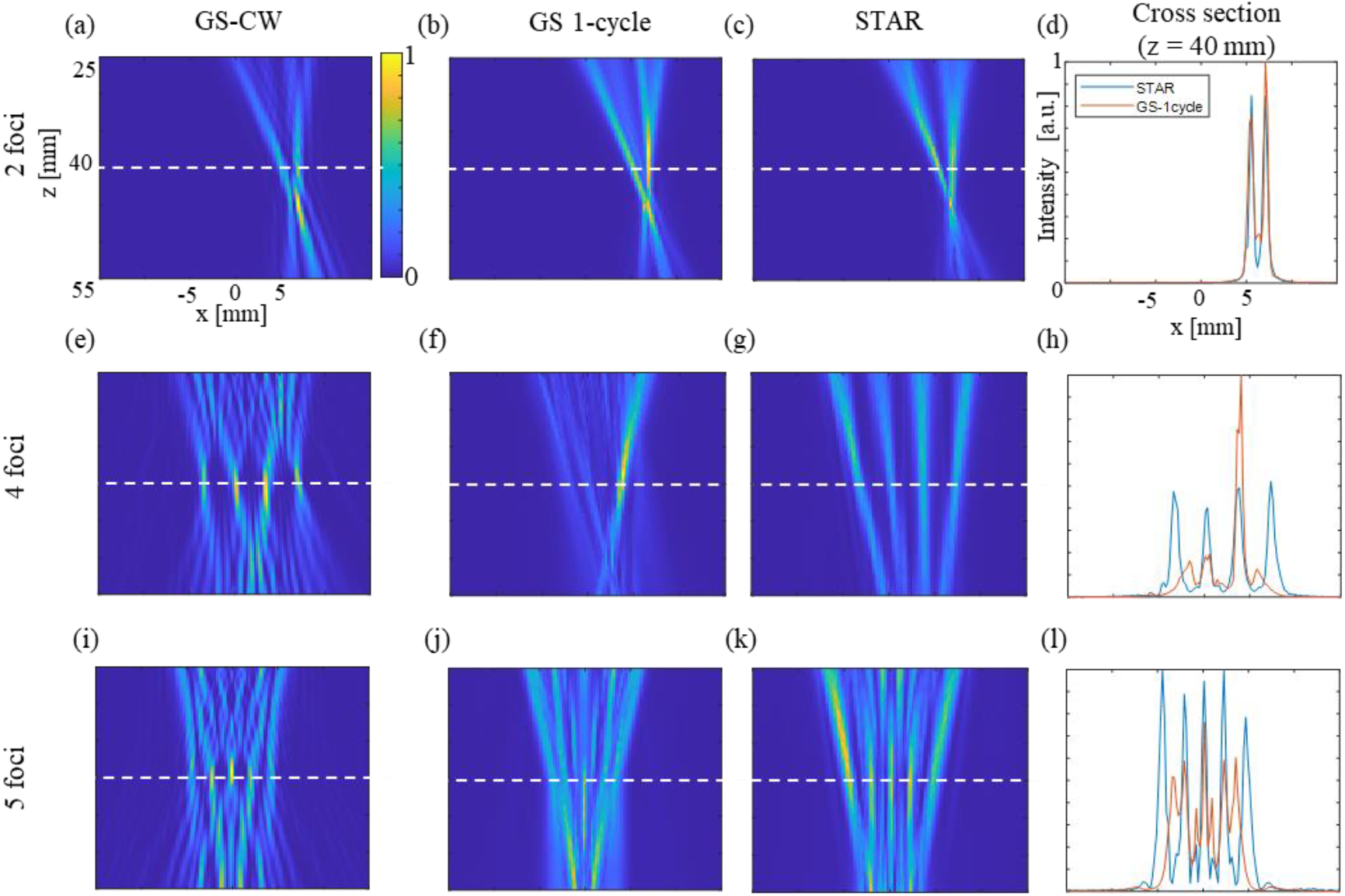
Comparison of GS and STAR algorithm performance across different multi-focus patterns: (a) XZ-plane intensity distribution for a two-foci pattern with a spacing of approximately 1.5 mm (7 pitches) and a 6 mm lateral offset, obtained using GS with CW excitation. (b) XZ-plane intensity distribution for the same pattern using GS with single-cycle excitation. (c) XZ-plane intensity distribution using STAR with single-cycle excitation. (d) Cross-section comparisons at the target depth of 40 mm for the two-foci pattern. (e) XZ-plane intensity distribution for a four-foci pattern with a spacing of approximately 3.5 mm (16 pitches) and a 3.5 mm lateral offset, obtained using GS with CW excitation. (f) XZ-plane intensity distribution for the same pattern using GS with single-cycle excitation. (g) XZ-plane intensity distribution using STAR with single-cycle excitation. (h) Cross-section comparisons at the target depth of 40 mm for the four-foci pattern. (i) XZ-plane intensity distribution for a five-foci pattern symmetrically arranged around the lateral axis, with a spacing of approximately 2.2 mm (10 pitches) between each pair, obtained using GS with CW excitation. (j) XZ-plane intensity distribution for the same pattern using GS with single-cycle excitation. (k) XZ-plane intensity distribution using STAR with single-cycle excitation. (l) Cross-section comparisons at the target depth of 40 mm for the five-foci pattern.

However, the STAR algorithm achieved a more balanced energy distribution between the two foci and produced a shallower null region between them—half as deep compared to the GS output (Fig. 5(d)). While the improvements provided by STAR for two-foci patterns with single-cycle excitation were relatively modest, since GS already performed well, STAR still achieved slightly better results. For instance, the correlation with the desired pattern was 0.869 for STAR compared to 0.861 for GS, reflecting an improvement of approximately 1%.

As the number of foci increased, the performance differences became more noticeable. The second pattern consisted of four foci spaced roughly 3.5 mm apart, with an offset of approximately 3.5 mm from the center of the lateral axis. In contrast, STAR successfully generated four nearly equal foci at the target depth (Fig. 5(g)). While the total energy at the target depth was only slightly higher in the STAR case, the energy distribution was more uniform between the foci (Fig. 5(h)). This improvement is also reflected in the correlation values, with STAR achieving 0.80 compared to 0.64 for GS. Additionally, as the number of foci increased, the accuracy of the GS output decreased relative to STAR. Specifically, positional errors became more pronounced—for example, in this case, the leftmost and rightmost foci were misplaced in the GS single-cycle output. The third pattern involved five foci symmetrically arranged around the lateral axis, with a spacing of approximately 2.2 mm (10 pitches) between each pair. Similar to previous cases, GS performed well for CW excitation (Fig. 5(i)) but failed to generate five distinct foci under single-cycle excitation (Fig. 5(j)). The foci were not well-defined, and the null regions between them were too high. However, STAR successfully produced the pattern, with accurate focal positions and minimal variation in peak heights (Fig. 5(k)). Notably, GS struggled with foci located further from the center of the lateral axis. While the two foci adjacent to the central one were positioned correctly, the outermost two were shifted closer to the center (Fig. 5(l)). Across all three patterns, and in approximately 95% of the tested patterns, the correlation between STAR results and the desired patterns was consistently higher than that of GS results. For two or more foci, GS achieved the highest correlation under CW excitation, as expected. However, STAR consistently outperformed GS in single-cycle excitation, particularly as the number of foci increased, ensuring better energy distribution, positional accuracy, and peak uniformity.

After evaluating the success of the STAR algorithm with various patterns at a depth of 40 mm and a frequency of 4.5 MHz, the scope of the research was expanded to include different target depths and frequencies. In the STAR algorithm, both parameters are defined as input values, similar to the GS algorithm. For this comparison, several patterns were tested. The presented example (Fig. 6) focuses on a 3-foci pattern symmetrically distributed around the center of the lateral axis, with a spacing of 12 pitches (∼2.6 mm) between each pair of foci. Both algorithms were tasked with generating this pattern at different target depths (30 mm, 40 mm, and 50 mm are displayed in Fig. 6).

**Fig. 6.**
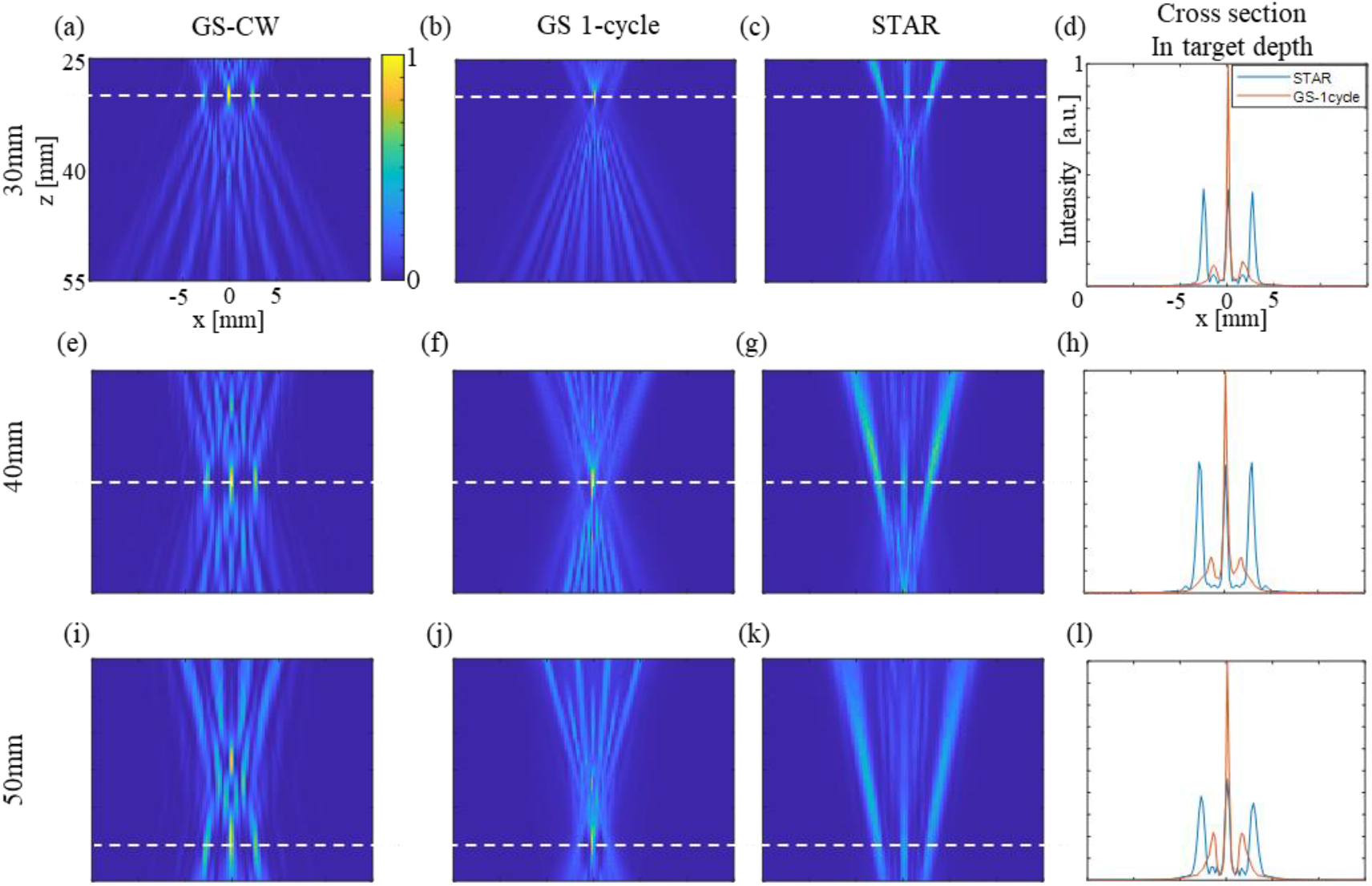
Comparison of GS and STAR algorithm performance for a 3-foci pattern with 12-pitch spacing (∼2.6 mm) between each pair of foci at different target depths. (a) XZ-plane intensity distribution at 30 mm depth using GS with CW excitation. (b) XZ-plane intensity distribution at 30 mm depth using GS with single-cycle excitation. (c) XZ-plane intensity distribution at 30 mm depth using STAR with single-cycle excitation. (d) Cross-section comparisons at 30 mm depth. (e) XZ-plane intensity distribution at 40 mm depth using GS with CW excitation. (f) XZ-plane intensity distribution at 40 mm depth using GS with single-cycle excitation. (g) XZ-plane intensity distribution at 40 mm depth using STAR with single-cycle excitation. (h) Cross-section comparisons at 40 mm depth. (i) XZ-plane intensity distribution at 50 mm depth using GS with CW excitation. (j) XZ-plane intensity distribution at 50 mm depth using GS with single-cycle excitation. (k) XZ-plane intensity distribution at 50 mm depth using STAR with single-cycle excitation. (l) Cross-section comparisons at 50 mm depth.

The GS algorithm performs well under CW excitation at all target depths, achieving a correlation of over 0.9 in all cases (Fig. 6(a), 6(e), 6(i)). The resulting patterns exhibit foci at the exact desired locations, with three distinct foci of slightly varying heights due to the apodization constraint, which was set to 1 for all elements. Allowing flexibility in apodization would equalize the foci heights and improve correlation, bringing it closer to 1, but would result in energy loss. However, with single-cycle excitation, the GS algorithm shows significant degradation in performance (Fig. 6(b), 6(f), 6(j)). The typical pattern consists of a high central focus on the lateral axis with two smaller, smeared foci. These smaller foci are not at the desired locations and are 5 to 10 times lower in height than the central focus (Fig. 6(d), 6(h), and 6(l)). This degradation is also evident in the correlation values, which are 0.55, 0.69, and 0.62 for target depths of 30 mm, 40 mm, and 50 mm, respectively. In contrast, the STAR algorithm performs consistently well across all examined depths. The 3-foci pattern is accurately generated, with the foci located at the intended positions and of equal heights (Fig. 6(c), 6(g), 6(k)). The correlation values are also much higher (0.82, 0.82, and 0.77). Although the two off-center foci exhibit slightly greater widths than the central focus, the STAR output distributes energy more evenly among the three foci compared to the GS output. While the GS single-cycle propagation results exhibit a higher central focus, the STAR output is superior in achieving equal focus heights and distributing energy more evenly among the three foci, which is a critical aspect of the desired pattern.

In addition to testing different depths, the algorithms were also evaluated at various frequencies around the transducer’s center frequency. Fig. 7 presents results for the same 3-foci pattern at a depth of 40 mm with frequencies of 3.5 MHz, 4.5 MHz, and 5.5 MHz. Across all tested frequencies, the STAR algorithm demonstrated superior performance compared to the GS algorithm, particularly in achieving uniform foci and consistent focus heights. The STAR algorithm achieved significantly higher correlation values (0.72, 0.82, and 0.76) compared to GS (0.54, 0.69, and 0.54) at frequencies of 3.5 MHz, 4.5 MHz, and 5.5 MHz, respectively. The GS algorithm consistently produced a higher central focus, accompanied by two smaller, less distinct foci that were mispositioned and asymmetrically smeared. The best results for uniformity and focus height were achieved at 4.5 MHz, likely due to it being the transducer’s center frequency. In general, as the frequency decreases, the average width of the foci increases, which is consistent with the expected degradation in lateral resolution at lower frequencies. This trend was observed in both GS and STAR algorithm outputs, as expected.

**Fig. 7.**
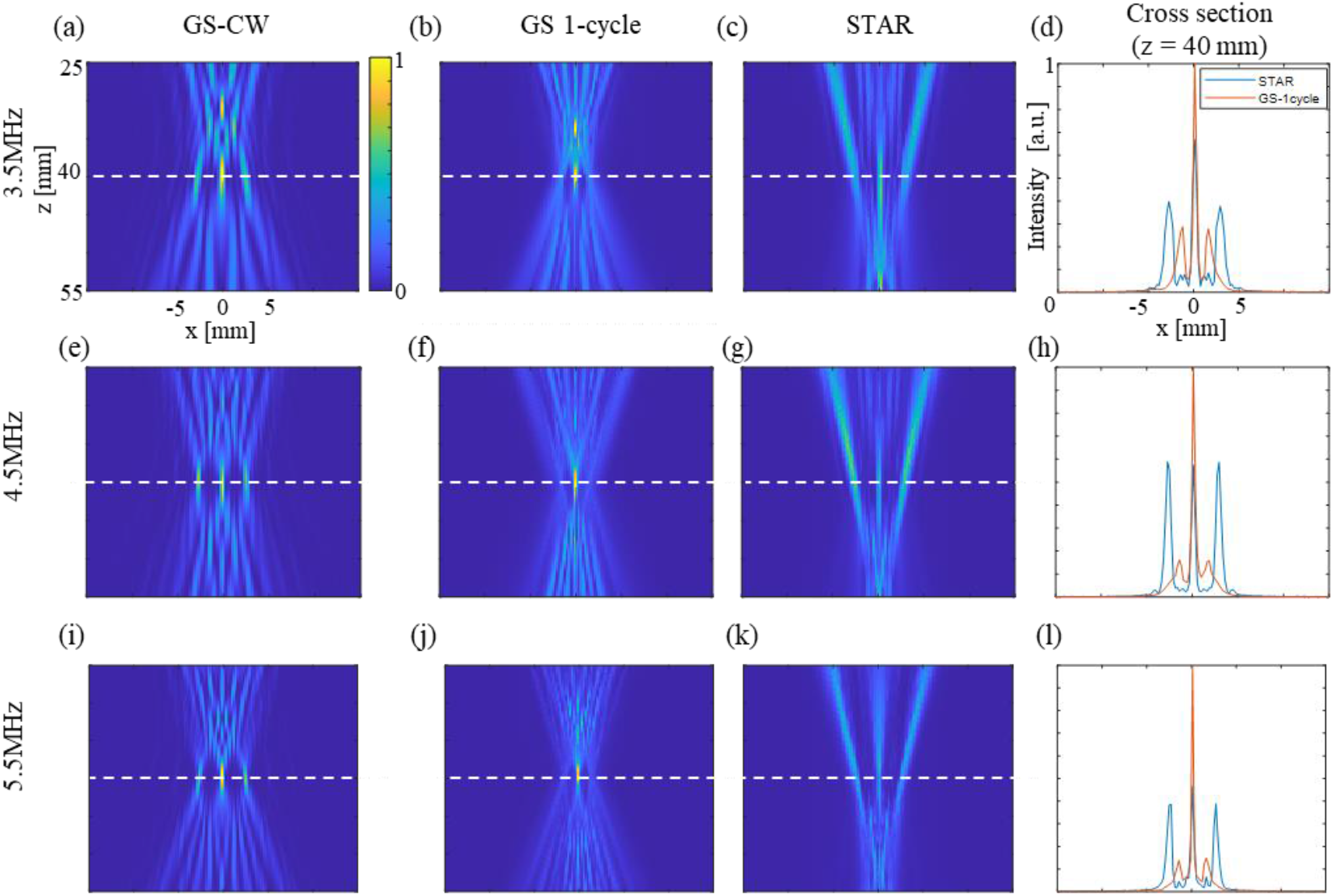
Comparison of GS and STAR algorithm performance for a 3-foci pattern with 12-pitch spacing (∼2.6 mm) between each pair of foci at 40 mm depth across different frequencies. (a) XZ-plane intensity distribution at 3.5 MHz using GS with CW excitation. (b) XZ-plane intensity distribution at 3.5 MHz using GS with single-cycle excitation. (c) XZ-plane intensity distribution at 3.5 MHz using STAR with single-cycle excitation. (d) Cross-section comparisons at 3.5 MHz. (e) XZ-plane intensity distribution at 4.5 MHz using GS with CW excitation. (f) XZ-plane intensity distribution at 4.5 MHz using GS with single-cycle excitation. (g) XZ-plane intensity distribution at 4.5 MHz using STAR with single-cycle excitation. (h) Cross-section comparisons at 4.5 MHz. (i) XZ-plane intensity distribution at 5.5 MHz using GS with CW excitation. (j) XZ-plane intensity distribution at 5.5 MHz using GS with single-cycle excitation. (k) XZ-plane intensity distribution at 5.5 MHz using STAR with single-cycle excitation.(l) Cross-section comparisons at 5.5 MHz.

### E. Resolution Analysis: Minimum Achievable Foci Spacing with GS and STAR

To evaluate the resolution improvement achieved by the STAR algorithm, it was of interest to determine whether STAR could generate patterns with closer foci compared to the GS algorithm. The process was as follows: the algorithms were tasked with generating two foci at progressively smaller spacings, aiming to reduce the spacing until the foci could no longer be distinguished (Fig. 8). The results indicate that both algorithms fail to produce two distinguishable foci at a spacing of 2 pitches and struggle at 3 pitches, although the STAR algorithm visually performs better in this case. However, while the GS algorithm fails to generate two distinct foci at a spacing of 4 pitches, the STAR algorithm successfully resolves them. Foci are considered distinguishable if the intensity between them drops to less than half of the maximum intensity of the smaller focus. Under the parameters of this study, the minimum achievable spacing was improved from 1.09 mm with the GS algorithm to 0.872 mm with the STAR algorithm for single-cycle excitation. Additionally, for larger distances, both algorithms successfully generated distinct foci.

**Fig. 8.**
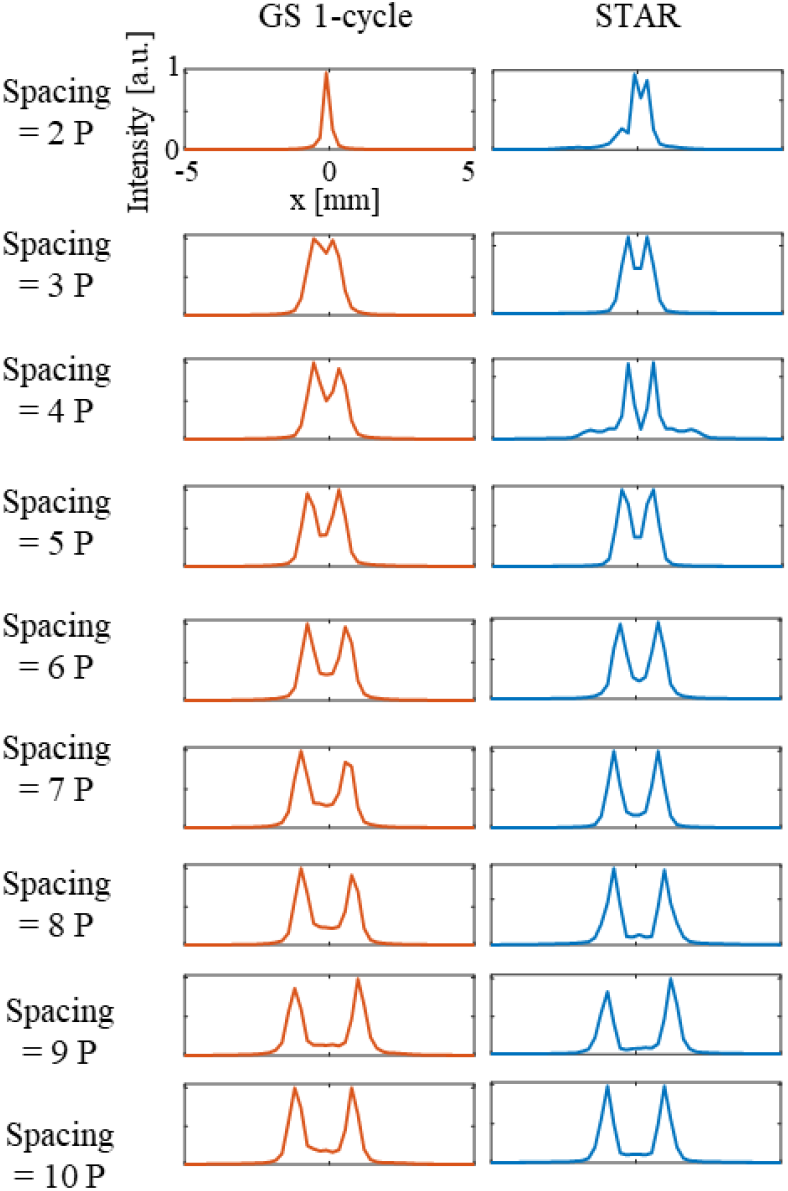
Resolution test-Comparison of GS and STAR algorithm performance in generating closely spaced. Cross-sections at the target depth for progressively increasing foci spacings. The left column shows the cross-section obtained from GS with single-cycle excitation, while the right column shows the cross-section from STAR with single-cycle excitation.

### F. Effect of Number of Foci on Pressure Intensity and Uniformity

When increasing the number of foci, a decrease in the average height of the foci is expected due to the redistribution of finite acoustic energy among a larger number of focal points. This behaviour is confirmed in the STAR algorithm outputs (Fig. 9(a)). The algorithm was tested across hundreds of patterns with varying numbers of foci and different spacings between adjacent foci. For each configuration, the pressure intensity at each focus was measured, and the mean pressure intensity was calculated. The results demonstrate a clear decrease in mean pressure as the number of foci increases.

**Fig. 9.**
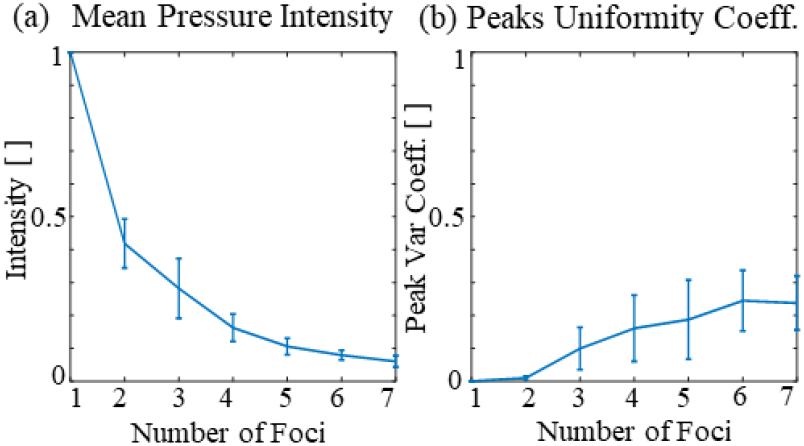
Effect of increasing the number of foci. (a) Mean pressure intensity at each focus as a function of the number of foci, measured across various patterns with varying numbers of foci and different spacings between each pair of foci. (b) Mean Peaks Uniformity Coefficient as a function of the number of foci, measured across the same patterns.

Additionally, the uniformity of focal peaks was analyzed using the Peaks Uniformity Coefficient, which was calculated for the same dataset (Fig. 9(b)). This parameter correlates with the difficulty of generating the desired pattern, assuming all foci are of equal height. The results show that as the number of foci increases, the Peaks Uniformity Coefficient also increases, although it remains relatively low. This finding aligns with the observation that patterns with one or two foci are generated with high accuracy and uniformity, while patterns with a larger number of foci occasionally display unequal energy distribution. This challenge becomes more evident as the number of foci increases, as highlighted in Fig. 9(b) and supported by Fig.5.

### G. Experimental Validation of the STAR Algorithm

To validate the proposed algorithm, experimental measurements were conducted using the setup described in the Methods section (Fig. 10). Four patterns were tested experimentally. Each column in the figure corresponds to a different pattern and each row represents a specific algorithm or signal duration. It should be noted that both XZ planes from the GS single-cycle and STAR outputs for the same desired pattern were normalized to the same scale to ensure a fair comparison, as the emitted energy was identical in both cases due to the single-cycle excitation.

**Fig. 10.**
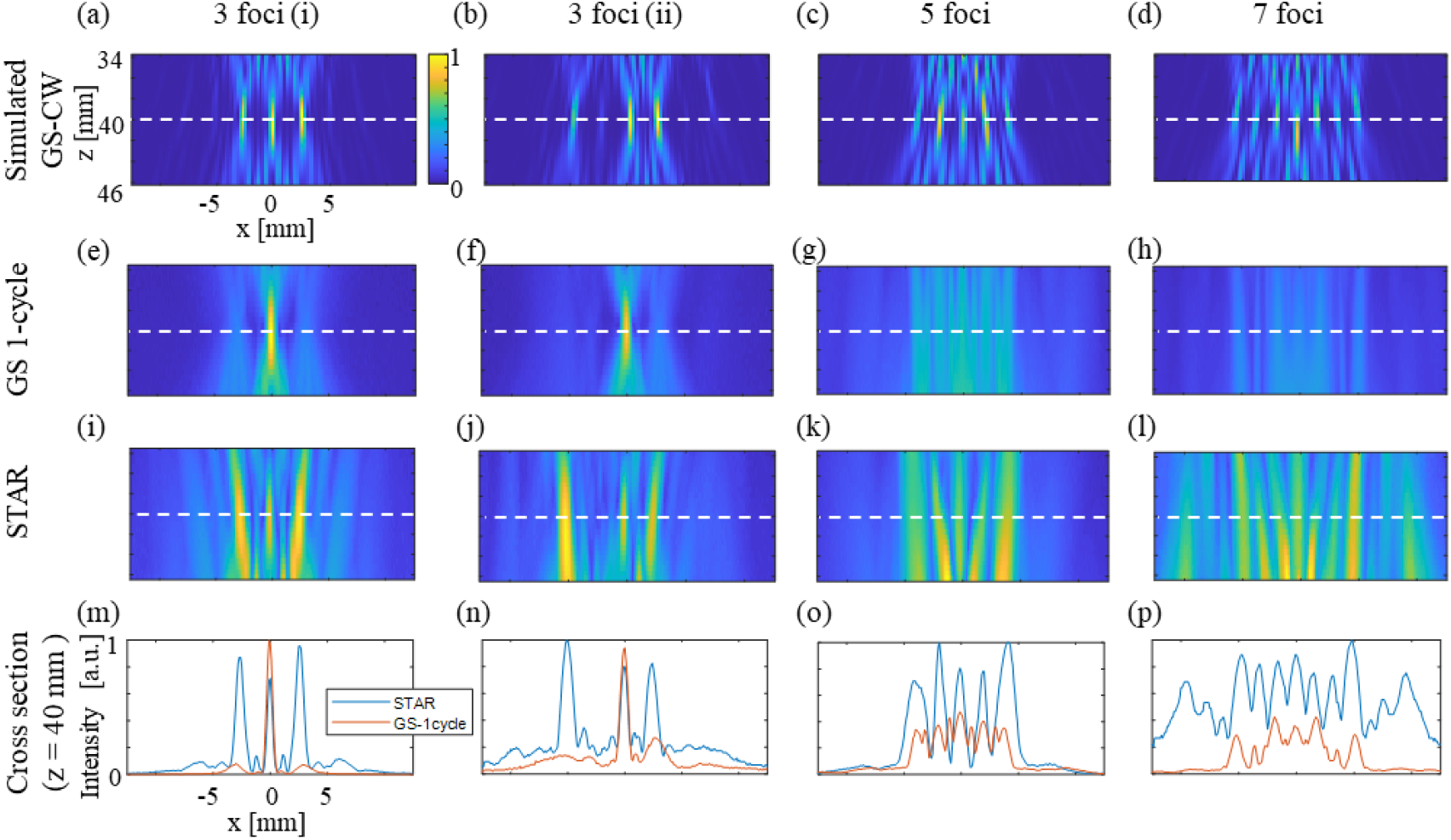
Experimental validation of GS and STAR algorithm performance for multi-focus patterns. Each column represents a different tested pattern, and each row corresponds to a different algorithm or excitation. The first pattern consists of three symmetrically a rranged foci with 12-pitch spacing. The second pattern features three asymmetrically spaced foci (11 and 23 pitches). The third pattern consists of five foci with 9-pitch spacing. The final pattern contains seven foci with 8-pitch spacing. (a–d) XZ-plane intensity distributions from simulations of continuous-wave (CW) propagation using GS-derived delays. (e–h) Measured acoustic fields from hydrophone recordings for XZ planes generated using GS-derived delays with single-cycle excitation. (i–l) Measured acoustic fields from hydrophone recordings for XZ planes generated using STAR-derived delays with single-cycle excitation. (m–p) Cross-section comparisons at the target depth of 40 mm for GS and STAR outputs.

The first pattern tested consisted of three foci symmetrically arranged around the central x-axis with a spacing of 12 pitches. The acoustic field measured for the GS single-cycle excitation (Fig. 10(e)) closely matches the simulated result from Fig. 6(f). Similarly, the acoustic field measured for the STAR output (Fig. 10(i)) correlates well with the simulation shown in Fig. 8(g). The measured cross-sections (Fig. 10(m)) also validate the simulation results from Fig. 6(h). Interestingly, in terms of energy distribution, the STAR algorithm demonstrated even better performance than predicted by the simulation, achieving higher relative focal intensities compared to the GS single-cycle cross-section. The GS output exhibited the typical pattern of one dominant focus with two weaker, smeared side foci. Conversely, the STAR output produced three foci with relatively close intensities, though not perfectly equal. The second pattern tested included three foci with unequal spacings (11 pitches and 23 pitches) arranged asymmetrically around the x-axis. The GS algorithm struggled to generate this pattern with single-cycle excitation, whereas the STAR algorithm demonstrated significantly superior performance (Fig. 10(n)). The difference in performance was even more evident in the third pattern, which consisted of five foci with equal spacing (9 pitches). The acoustic field generated from the GS output for this pattern did not adequately reproduce the desired configuration, making it challenging to identify the original pattern (Fig. 10(g)). In contrast, the STAR algorithm successfully generated five foci with nearly equal intensities at the specified positions. The higher relative intensity of the STAR-generated foci, compared to GS, further highlights its ability to concentrate energy more effectively at the target depth (Fig. 10(o)). Additionally, it was consistently observed that foci located further from the center of the lateral axis tended to be broader. This behavior was apparent across all patterns and was particularly evident in Fig. 10(o). The final pattern consisted of seven foci with a spacing of 8 pitches between each pair. Both algorithms exhibited limitations in generating this pattern effectively in the experiments (Fig. 10(p)). For the GS single-cycle output, the foci were weak and indistinct. The STAR algorithm produced high side lobes in addition to the desired pattern. Although the foci generated by the STAR algorithm were not completely distinct, they were more distinguishable compared to the GS output.

## IV Discussion

This study introduces the STAR algorithm as a generalized method for beam shaping of ultra-short pulses across wave-based technologies, with its capabilities demonstrated in medical ultrasound. By formulating the optimization in the time domain, STAR enables explicit control over excitation duration, including single-cycle pulses. A critical element of the implementation is the integration of the GASM, which provides efficient FFT-based propagation and back-propagation between parallel planes, thereby supporting iterative optimization.

The accuracy of the GASM in modeling acoustic propagation was validated by strong agreement with the k-Wave toolbox across multiple patterns and depths. In addition, by restricting computations to specific planes rather than entire volumetric grids, the GASM achieved propagation times several orders of magnitude faster, establishing it as an efficient and reliable propagation engine for STAR.

For simple patterns, such as a single focus, the STAR algorithm converged to the GS solution, consistent with the analytical geometrical solution. Since this solution is considered ideal, it provided a reliable validation step for both GS and STAR implementations. The advantages of STAR with single-cycle excitation became particularly evident as pattern complexity increased. For multi-foci patterns, STAR achieved higher correlation with the desired patterns, greater energy concentration, more uniform focus heights, and improved positional accuracy compared to GS, which often produced smeared or uneven intensities. These improvements were observed across a wide range of configurations, including patterns with increasing numbers of foci, varying lateral spacings, and asymmetric arrangements. Notably, the minimum distinguishable spacing improved from 1.09 mm with GS to 0.872 mm with STAR, demonstrating its capacity to resolve closely spaced features. Robust performance was further confirmed across depths (30–50 mm), excitation frequencies (3.5–5.5 MHz), and numbers of foci (2–7). The best uniformity was obtained at 4.5 MHz, consistent with the transducer’s center frequency, while lower frequencies produced broader foci in line with the expected degradation of lateral resolution. Importantly, experimental hydrophone measurements validated the simulation results, with STAR consistently reproducing target patterns more accurately than GS.

While STAR consistently outperformed GS, challenges remained for highly complex patterns, such as seven foci with close spacing. Even in these cases, however, STAR showed a clear advantage in focus distinguishability and energy concentration. Incorporating additional stopping criteria, such as the Peaks Uniformity Coefficient, can further improve uniformity, though at the cost of reduced flexibility when focus heights vary. This metric was provided as an example, and alternative criteria could be adopted depending on the application. In practical tissue imaging, additional complexities may arise due to aberrations, which should be systematically investigated. Such effects could be mitigated either by integrating aberration-correction algorithms or by explicitly modeling the heterogeneous medium within the optimization process.

Time-domain processing has been applied to ultrasound imaging on the receive side [36], highlighting its value for beamforming. However, such developments remained focused on reception, whereas STAR uniquely addresses active beam shaping in transmission.

Future studies should extend STAR to complete imaging schemes that combine transmission and reception, thereby opening new opportunities for improved resolution and advanced ultrasoud imaging. Beyond ultrasound, the adaptability of the time-domain framework suggests potential applications in optics, radar, and photoacoustics, where precise shaping of ultra-short pulses is essential. Further work could also explore polychromatic excitations and diverse pulse shapes, leveraging STAR’s ability to handle arbitrary time-domain signals.

In conclusion, the STAR algorithm, integrated with the GASM, provides a versatile and efficient framework with the potential to significantly enhance ultrasound imaging and other wave-based applications.

## V Conclusion

This study introduced the STAR algorithm as a generalized beam-shaping method for ultra-short pulses, demonstrated in the field of medical ultrasound. By integrating STAR with the GASM, the proposed approach achieves precise multi-foci generation while maintaining high computational efficiency, particularly for single-cycle excitations. This framework addresses key limitations of phase-based methods, offering improved spatial accuracy, energy distribution, and adaptability. The findings have significant implications for biomedical ultrasound, particularly in applications requiring precise control over acoustic fields, such as high-resolution and high frame rate imaging.

## Appendix

Supplementary Figure is attached.

---

**Supplementary Fig. 1.**
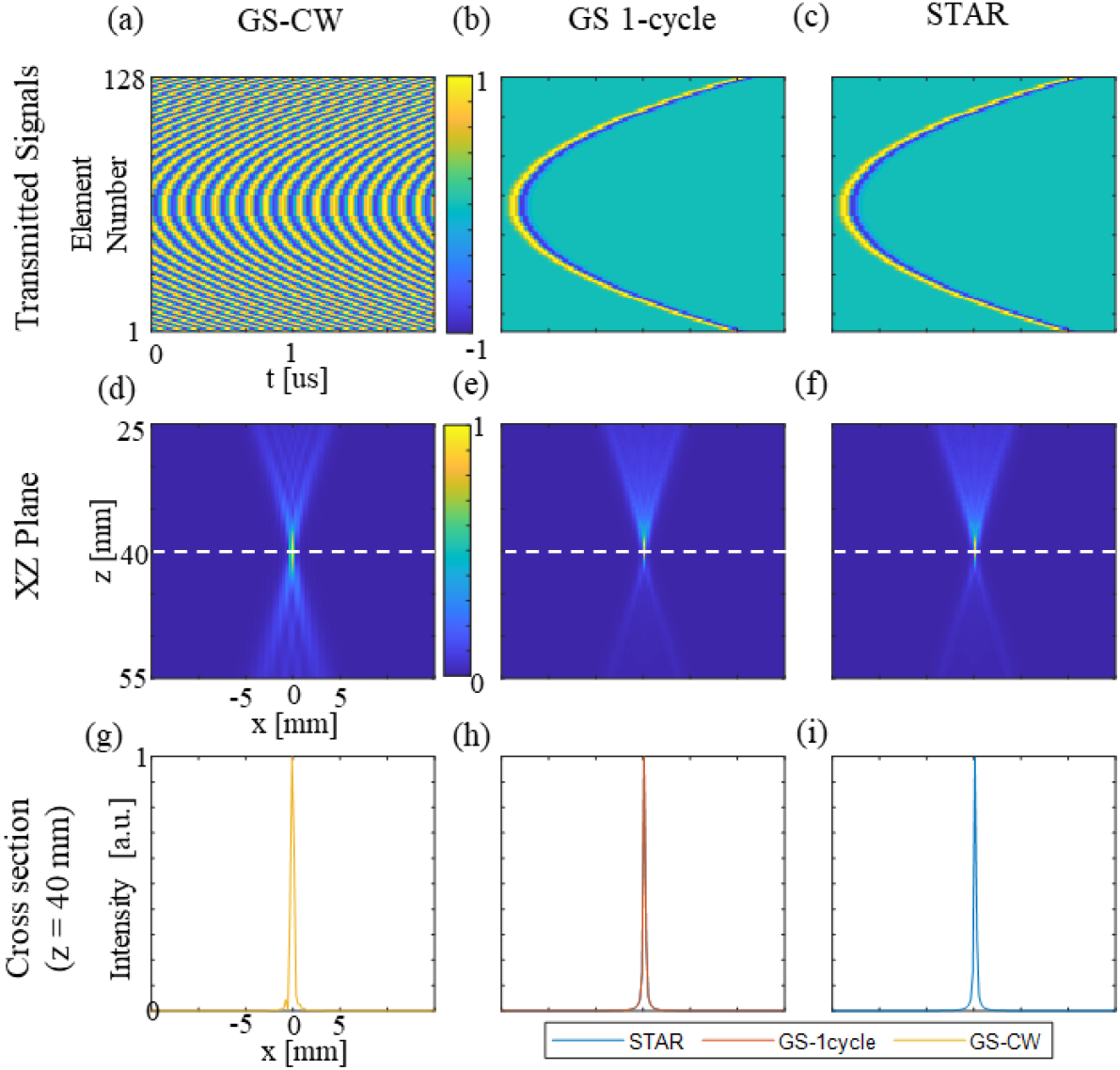
Validation of the GS and STAR algorithms for a single-focus pattern at a depth of 40 mm (4.46 MHz). (a) Wavefronts generated using the GS algorithm with an input pattern of a single focus at the center of the lateral axis. (b) Corresponding wavefronts for single-cycle excitation. (c) Wavefronts from the STAR algorithm. (d–f) XZ-plane intensity distributions from GS with CW excitation, GS with single-cycle excitation, and STAR, respectively. (g–i) Cross-section comparisons at 40 mm depth for the three cases.

